# A Unique Ribosome Signature Reveals Bacterial Translation Initiation Sites

**DOI:** 10.1101/095893

**Authors:** Adam Giess, Elvis Ndah, Veronique Jonckheere, Petra Van Damme, Eivind Valen

## Abstract

While methods for annotation of genes are increasingly reliable the exact identification of the translation initiation site remains a challenging problem. Since the N-termini of proteins often contain regulatory and targeting information developing a robust method for start site identification is crucial. Ribosome profiling reads show distinct patterns of read length distributions around translation initiation sites. These patterns are typically lost in standard ribosome profiling analysis pipelines, when reads from footprints are adjusted to determine the specific codon being translated. Using these unique signatures we build a model capable of predicting translation initiation sites and demonstrate its high accuracy using N-terminal proteomics. Applying this to prokaryotic samples, we re-annotate translation initiation sites and provide evidence of N-terminal truncations and elongations of annotated coding sequences. These re-annotations are supported by the presence of Shine-Dalgarno sequences, structural and sequence based features and N-terminal peptides. Finally, our model identifies 61 novel genes previously undiscovered in the genome.

---

Identification of translated open reading frames (ORFs) is a critical step towards annotation of genes and the understanding of a genome. In addition to providing functional information via the peptide sequence, regulatory and targeting information are often contained within protein N-termini^1,2^, this makes accurate identification of the beginning of ORFs essential. Whole genome ORF identification in prokaryotes is most commonly performed *in silico*, using a variety of sequence features, such as GC codon bias and motifs such as the ribosomal binding site or Shine-Dalgarno sequence^3,4,5^ in order to differentiate those ORFs that are thought to be functional from those that occur in the genome by chance. While these techniques are able to identify genomic regions containing ORFs with a high accuracy^5^, predicting translation initiation sites (TISs), and thus the exact beginning of a protein coding sequence (CDS), is substantially more challenging. This has led to the development of a number of *in silico* based TIS identification methods relying on a variety of sequence features^6-9^, which are typically post processing tools applied after initial ORF annotation in order to re-annotate the often erroneously predicted TIS.

High throughput proteogenomics has the potential to enable identification of protein N-termini, and by extension TISs, from an entire proteome. In practice however variation in protein expression levels, physical properties, MS-incompatibility and the occurrence of protein modifications limit the number of detectable protein N-termini^10,11^. In prokaryotes N-terminal proteomics typically captures the corresponding peptides of hundreds to the low thousands of genes^11^. For example, a recent study identified N-terminal peptides of 910 of the 4140 (22%) annotated genes in *Escherichia coli^12^.* Although falling short of providing full genome annotation, such datasets provide an effective means of experimental TIS validation.

Significantly higher coverage of TISs can be achived by using sequencing based technologies. By specifically focusing on ribosome protected fragments, ribosome profiling^13^ (ribo-seq) infers which parts of the transcriptome are actively undergoing translation. In this way, ribo-seq has been used to demonstrate translation of many RNAs and regions that were not thought to be associated with ribosomes^14-20^. Being able to identify translation on a transcriptome-wide scale has obvious application to ORF annotation and a number of methodologies have been developed for prediction of translated ORFs^17,19,21-23^. These methods rely on a number of features, like codon periodicity, read context and read lengths, in order to distinguish footprints indicative of translation from other, non-translating, footprints frequently observed in ribo-seq data. While extensive progress has been made on finding translated regions, delineating their exact boundaries has received less attention. Antibiotic treatment can be used to stall and capture footprints from the initiating ribosome^14,24,25^, but finding a suitable compound has been elusive in prokaryotes with only one dataset to date^26^.

Here we present a generally applicable method that does not depend on specialised chemical treatment, but can be take advantage of such data (Figure 1a). Using N-terminal proteomics we demonstrate its high accuracy and show that it is consistent with other features linked to translation initiation. Applying the model we predict numerous novel initiation sites in *Salmonella enterica* serovar Typhimurium.

**Figure 1:**
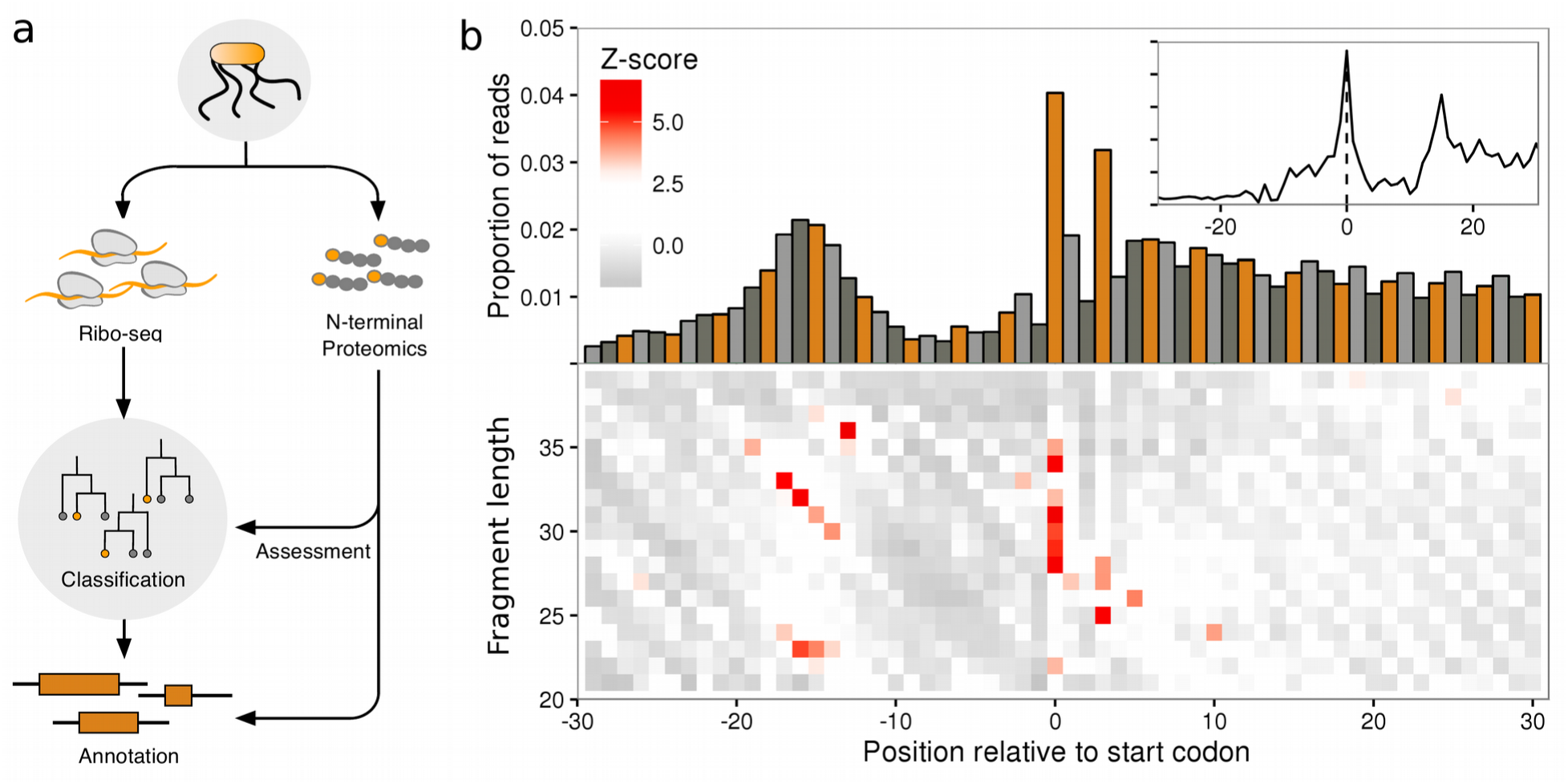
Translation initiation site classification with ribo-seq fragment length patterns. (**a**) Schematic representation of the classification strategy. (**b**) Ribo-seq meta profiles in windows around start codons for all annotated CDSs in the *S.* Typhimurium genome (monosome sample, n=4187), contributions from each gene are scaled to a sum of one. (**upper**) Proportion of 5’ ribo-seq read counts per nucleotide position, coloured by codon position. (**lower**) heatmaps of z-scores of 5’ ribo-seq read counts per fragment length. (**inset**) proportions of ribo-seq read counts per nucleotide position, after adjusting reads by fragment length offsets (see methods).

## RESULTS

### Translation Initiation Sites Carry a Unique Signature

To investigate whether ribo-seq could aid in the accurate delineation of translated ORFs we generated two ribo-seq libraries from monosome and polysome enriched fractions originating from *S.* Typhimurium. The similarities in the profiles of the two libraries (Supplementary Fig. S1e-f), taken with current literature reports of similarities in the translational properties of polysome and monosome fractions^27^, suggest that it is reasonable to consider these libraries sufficiently similar to serve as replicates for the purpose of initiation sites. The libraries were initially processed in a standard ribo-seq work-flow, where trimmed footprints were aligned to a reference genome, and then adjusted based on 5’ read profiles to determine the specific codon under active translation (Figure 1b, inset, Supplementary Fig. S1e-f, inset). When exploring the processed reads we discovered that, consistent with previous reports^26,28^, annotated start sites of ribosomes treated with chloramphenicol carry a unique signature around the initiating codon (Figure 1b, inset). Examining the unprocessed reads we observed that the pattern is a consequence of a specific distribution of fragment lengths (Figure 1b), information which is typically lost in pipelines that pre-process the read signal by adjusting reads (Figure 1b, inset). More specifically, heatmaps of 5’ read profiles indicate that the pattern consists of an enrichment of longer fragments (30-35 nucleotides(nt)) starting 14-19 nt upstream of the initiation codon (a diagonal pattern), but ending at the same location, 15 nt downstream of the initiation codon. A shorter set of fragments (23-24 nt) are enriched in the same region, but have different end points, 7-9 nt downstream of the initiation codon. And finally, a strong enrichment of 5’ ends of reads of length 28-35 nt can be observed exactly over the start codon itself (Figure 1b).

### Ribosome profiling enables accurate annotation of translation initiation sites

We trained a random forest model on TISs from the top 50% translated ORFs (see methods), to recognise the patterns in 5’ ribo-seq read lengths and sequence contexts in a -20 to +10 nt window around start codons. In addition we encoded information about the start codon position within the ORF and the read abundance upstream and downstream of the start sites. The model was then used to predict TISs from all in-frame cognate and near cognate (one edit distance from ATG) start codons around annotated genes in the *S.* Typhimurium genome. Predictions on the two samples were highly accurate with area under curve (AUC) values of 0.9958 and 0.9956 on independent validation sets for the monosome and polysome sample, respectively (parameter importance for the models is summarised in Supplementary Tables S1-2). In total 4610 (monosome) and 4601 (polysome) TISs were predicted in the two sets. From these, we constructed a high confidence set from predictions common to both replicates. In total this set contained 4272 predictions, representing an 86.50% agreement between the replicates. The discrepancies predominantly originate from genes with scarce translation. Of the high confidence TISs, 3853 matched annotated ORFs, 214 represented elongations and 205 truncations. Examples of predicted elongated, truncated and matching ORFs are shown in figure 2.

**Figure 2:**
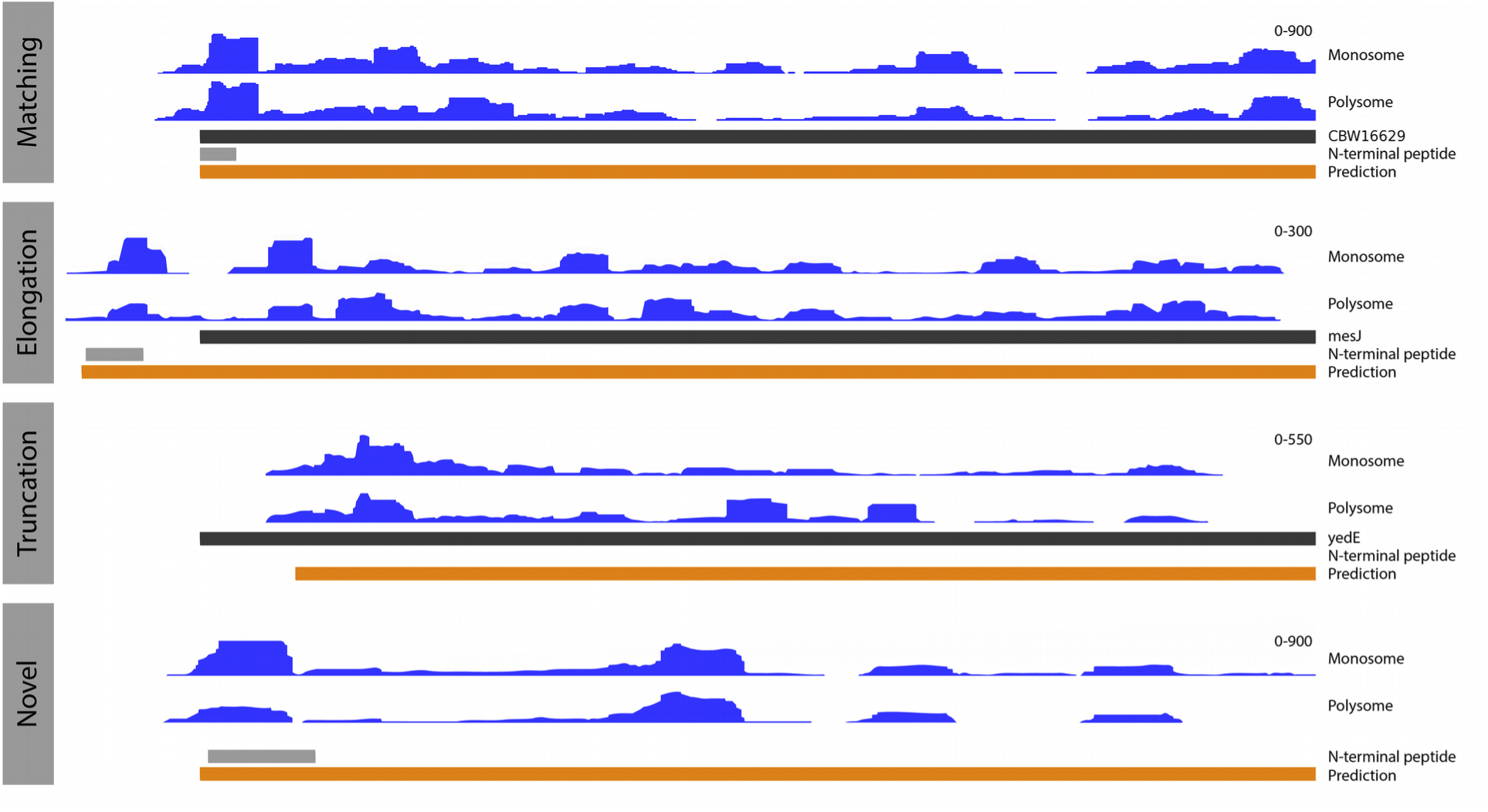
Examples of predicted translated ORFs. Showing genomic tracks of unadjusted ribo-seq read coverage in blue (y axis scale on the right hand side), annotated genes in black, predicted ORFs in orange and N-terminal peptides in grey. (**upper**) A predicted ORF in agreement with the annotated ORFs, supported by ribo-seq coverage and N-terminal evidence. (**middle upper**) A predicted elongation relative to the annotated ORF, with N-terminal evidence and ribo-seq coverage supporting the elongated prediction. (**middle lower**) a predicted truncation relative to the annotated ORF, with support from ribo-seq coverage. (**lower**) a novel predicted ORF, supported by N-terminal evidence and ribo-seq coverage.

As expected the predictions show the same codon usage distribution (Supplementary Fig. S2), and carry the same read distribution signature as the annotated sites (Supplementary Fig. S2). Consistent with annotated initiation sites an increase in ribosome protected fragments can be seen downstream versus upstream of the predicted TIS (Figure 3a). Furthermore, elongated ORFs exhibit a shift in ribo-seq density downstream relative to the annotated TIS, consistent with the predicted elongation. Conversely, truncated ORFs exhibit a shift in read density upstream relative to the annotated TIS and consistent with the predicted truncation.

**Figure 3:**
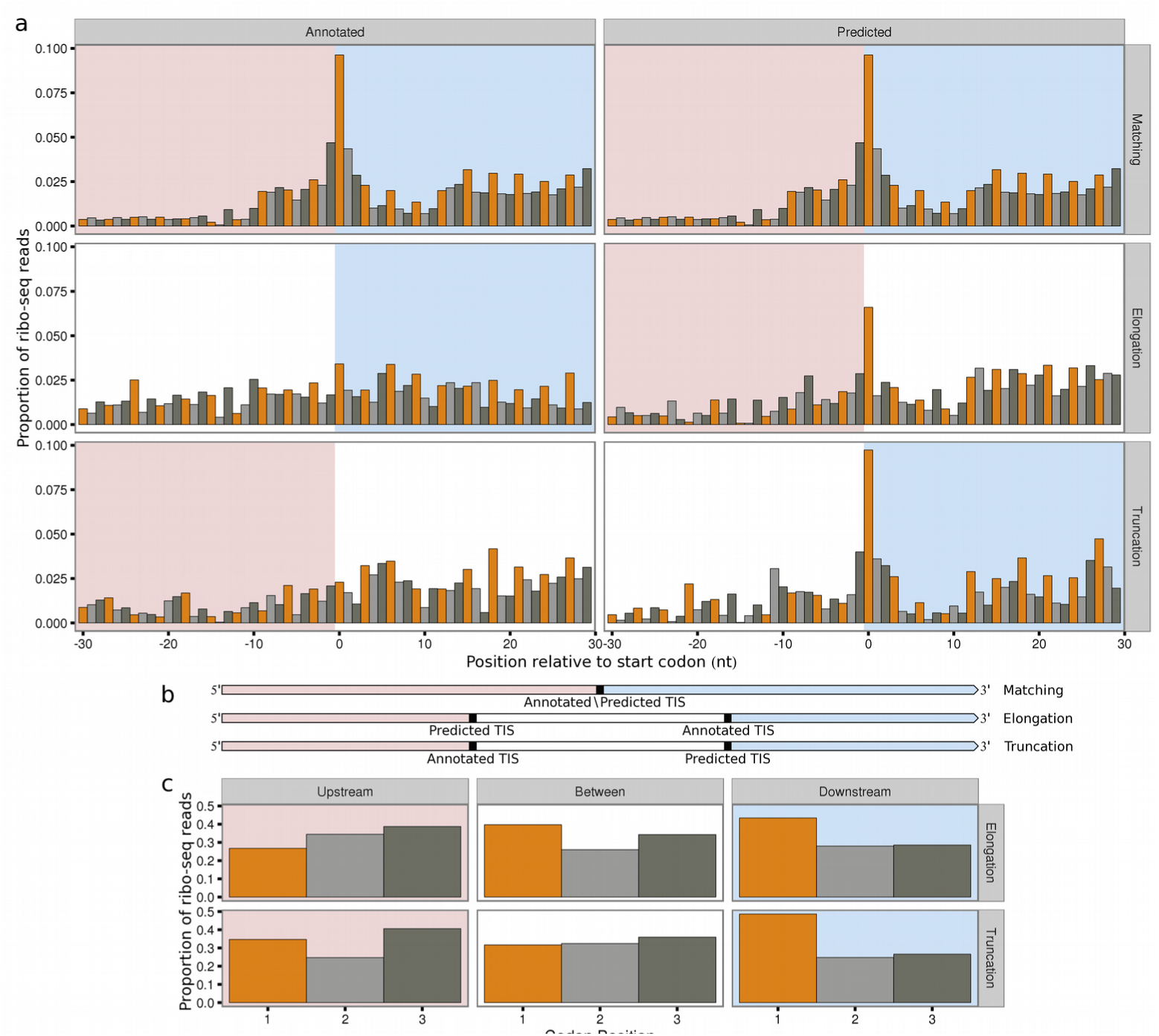
Ribo-seq reads and periodicity are consistent with re-annotated translation initiation sites. Bar colour indicates codon position. Downstream regions are highlighted in pink, upstream regions are highlighted in light blue. (**a**) Meta plots showing the proportion of scaled ribo-seq reads in relation to annotated or predicted translation initiation sites, for ORFs matching annotated genes (n=3853), predicted elongations (n=214) or predicted truncations (n=205). Contributions from each gene are scaled to a sum of one. Annotated TISs show increased ribo-seq density upstream (elongations), or downstream of start codons (truncations). (**b**) Transcript models. (**c**) Bar plots showing the sum of proportions of scaled ribo-seq read counts in each codon position. For truncations regions are 30 nt upstream of the annotated TIS, between the annotated and predicted TIS and 30 nt downstream of the predicted TIS. For elongations regions are 30 nt upstream of the predicted TIS, between the predicted and annotated TIS, and 30 nt downstream of annotated TIS. Three nt periodicity does not occur upstream of predicted TISs (truncations), but does occur upstream of annotated TISs (elongations).

To further assess the predictions we compared the newly predicted TISs with the previously, potentially erroneously, annotated TIS. A highly significant sequence feature of translation initiation sites is the Shine-Dalgarno (SD) sequence which facilitate translation initiation in prokaryotes^29^.The consensus sequence GGAGG is located approximately 10 nt upstream of the start codon^30^.The predicted initiation sites show clear evidence of SD sequences centred 9-10 nt upstream of the start codon (Figure 4a). Strikingly the annotated TISs, in these same genes where our model has predicted novel sites, show an absence of the SD sequence (Figure 4a). Since our model evaluates sequence context it is unsurprising that the predictions carry this signature, but the absence of these motifs around previously annotated start codons is notable.

**Figure 4:**
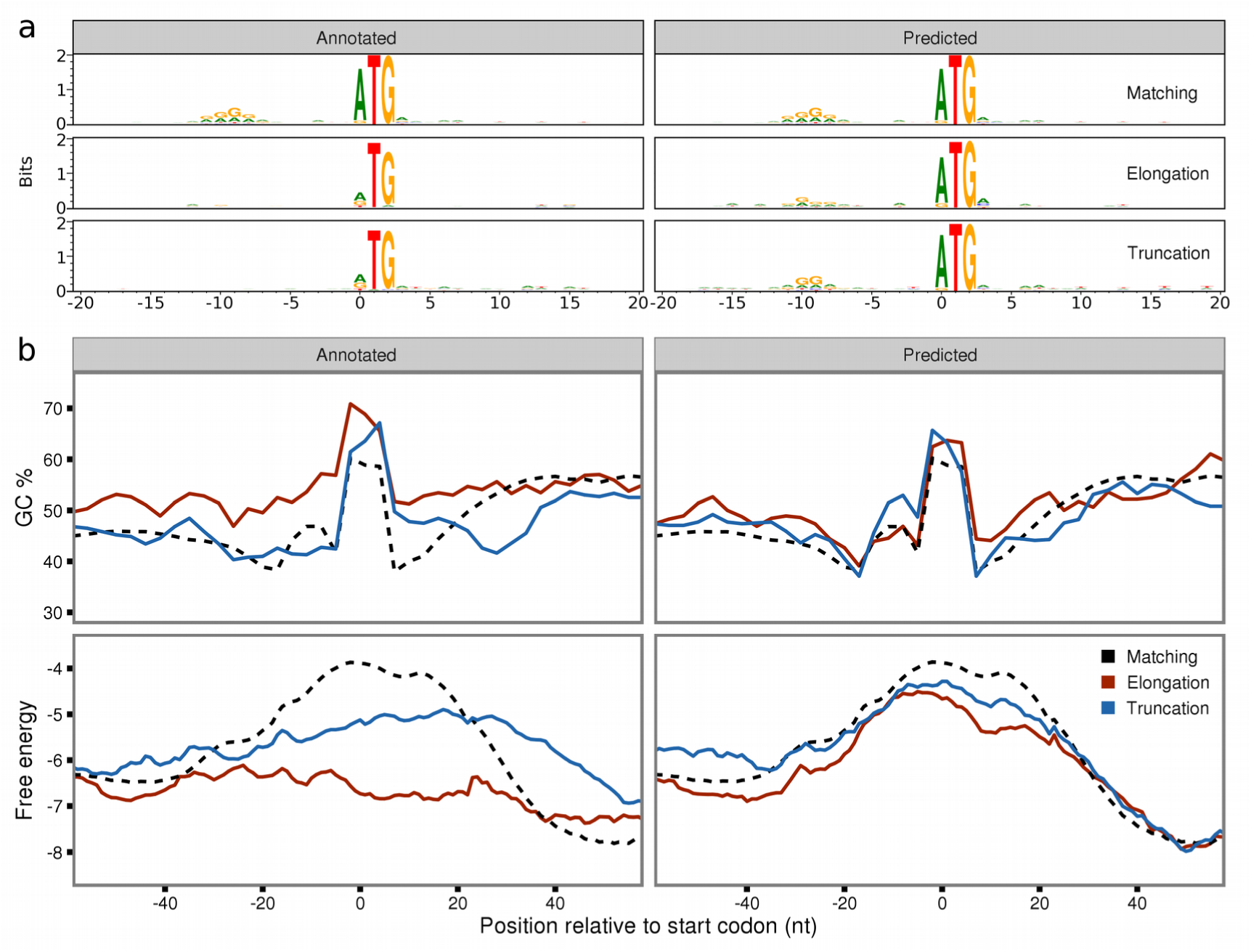
Sequence and structure features support re-annotation of translation initiation sites. (**a**) Sequence motifs relative to annotated or predicted translation initiation sites in the same genes. ‘Matching’ (n=3853) are identical, while predicted elongations (n=214) and truncations (n=205) have stronger SD sequences than their annotated counterparts. (**b**) Meta-profiles relative to annotated or predicted translation initiation sites, with lines representing ORFs matching annotated genes (dashed black), predicted elongations (red) and predicted truncations (blue). (**upper**) Mean GC content at third codon positions, averaged over 9nt sliding windows. Predicted TIS match the expected profile, showing an increase in GC content immediately after the start codon. Whereas predicted elongations and truncations show shifts down or upstream in annotated TIS. (**lower**) Meta-profiles of mean free energy averaged in 39 nt sliding windows. Peaks of low secondary structure potential, expected to occur over start codons, are centred over predicted TIS, but are clearly shifted down or upstream of annotated TIS, in predicted elongations and truncations.

Besides the presence of SD sequences, the guanine-cytosine (GC) content is commonly used to identify CDSs in prokaryotes. The overall GC content of a genome or genomic region is often highly optimised. In coding regions this optimisation can be achieved via synonymous substitutions, predominantly at third codon positions^31^, leading to a pronounced bias in the GC content of third nucleotide positions in coding regions compared to the rest of the genome. Interestingly at annotated sites, predicted elongations exhibit an increase in GC content upstream of the annotated start codon consistent with the location of the predicted site, conversely predicted truncations show a decrease downstream of the annotated start codon. In contrast, at predicted sites both predicted elongations and truncations fit closely to the expected distribution (Figure 4b upper).

Another significant feature of prokaryotic translation initiation is the absence of intrinsic structure in the region around the start codon enabling easier access for ribosomes to bind^32^. We therefore calculated the average free energy over all predicted sites and compared them to the previous annotation in the same genes. Consistent with GC content patterns, the annotated sites display a lower propensity to form secondary structure upstream of the start codon in elongated ORFs, and downstream of the start codon in truncated ORFs. In the predicted sites these less-structured regions can clearly be observed directly over the start codon, highly indicative of true initiation sites (Figure 4b lower).

Ribosomes translocate along mRNAs three nucleotides at a time, corresponding to one codon and amino acid (aa). Consequently, reads originating from bona fide translated regions also exhibit a three nucleotide periodicity in adjusted read counts, with a bias towards mapping to the first nucleotide in each codon^21^. At initiation sites read distribution therefore switches from a random distribution upstream to a periodic, biased distribution downstream. Comparing the density of reads falling into each of the three codon positions, in elongated ORFs we observe increased read density at the first nucleotide position upstream of annotated, but not predicted TISs. Similarly at truncated ORFs we see a decrease in the density of reads at the first nucleotide position downstream of the annotated TIS but not the predicted TIS (Figure 3c).

Taken together the patterns in read distribution, SD motifs, GC bias, unstructured regions and triplicate periodicity, provide clear and consistent support that the TISs which we reannotate, show on average, a higher agreement with features indicative of canonically translated prokaryotic ORFs, than their corresponding previously annotated counterparts.

### N-terminal proteomics confirms predicted sites

In order to experimentally validate the accuracy of the predictions positional proteomics analyses enriching for protein N-termini were performed. Blocked N-termini were identified at 1040 *S.* Typhimurium ORFs, from which a high confidence subset of Nt-formylated Met-starting N-termini was selected (see methods) and used for assessing the accuracy of the model. In total 114 high confidence N-termini were identified supporting 102 annotated CDSs, three N-terminal CDS elongations, and nine N-terminal CDS truncations. Because genomic positions with N-terminal peptide support were excluded from the set used to train the random forest model, these high confidence TIS positions can be used to determine the accuracy of the predictions. Of the 102 N-terminally supported annotated genes, 96 were predicted by the model. Furthermore two of the elongations, and four of the truncations were captured (Supplementary Table 3). Assuming that none of these genes have multiple initiation sites the sensitivity of the model can be estimated to be 0.9444, the specificity to be 0.9991 and the positive predicted value to be 0.9444.

The remaining set of blocked N-termini supported a further 668 annotated CDSs, 50 suggested CDS elongations and 310 suggested CDS truncations. In addition, we found peptides matching the predicted start positions from three distinct novel regions (defined as ORFs at least 300 nt in length, in regions that were not overlapping with annotated genes or regions at least 999 bp upstream of annotated genes). Comparing the predictions to the wider blocked N-termini set we find support for 648 predictions that match annotated TISs, 22 predicted elongations, 23 predicted truncated and three novel regions (Supplementary Table S4).

### Translation initiation sites are predicted at novel genomic regions

In order to discover potential novel translated ORFs we applied our prediction models to look for TISs in genomic regions outside annotated ORFs. Novel ORFs that were similar in size to known CDSs (> 100aa) and with ribo-seq coverage along a high proportion of the ORF (>75 % coverage, see methods) were considered candidate translated novel ORFs. Of the 219 (monosome) and 193 (polysome) ORFs under consideration, 104 and 115 novel translated ORFs were predicted respectively. 61 of these novel translated ORFs were common to both replicates (38.61% agreement) and used as a high confidence set of novel predictions. Unlike the annotated genes, these novel ORFs are not previously confirmed as translated regions and most had significantly lower read density (mean FPKM of 8) than annotated genes (mean FPKM of 126). The higher discrepancy between the two replicates is mainly a consequence of low-abundance start sites that did not pass the threshold in either of the replicates.

Read density plots over the novel ORFs revealed features consistent with protein coding regions, but with higher variance due to the low number of ORFs. Specifically, GC content increases downstream of the initiation codon, the regions around the initiation codon have less intrinsic structure potential and Shine-Dalgarno sequences are present upstream (Supplementary Fig. S3). Additionally, three of the predicted novel translated ORFs were supported by N-terminally enriched peptide evidence (a representative example is shown in Figure 2). A further 22 showed high similarity to known protein sequences, four of which contained functional protein domains (Supplementary Table S5).

### Tetracycline treated samples improve classifier accuracy

While reads isolated from elongating ribosomes provide sufficient information to predict the majority of translation start sites we set out to explore the full potential of our classifier in combination with publicly available data from initiating ribosomes. A recent study on *E. coli*^12^ demonstrated the use of tetracycline as a translation inhibitor to enrich for footprints from initiating ribosomes in prokaryotes. The tetracycline datasets show the pattern that we expect to see from initiating ribosomes as a range of read lengths starting 28-14nt upstream of the initiation codon (5’ data), but ending at the same positions 14-15nt downstream of the initiation codon (3’ data). An additional pattern of shorter fragment lengths can also be observed starting 26-18nt upstream, and ending 2nt downstream of the initiation codon (Supplementary Fig. S1.a,b,g,h).

We trained separate classifiers on chloramphenicol (elongating) and tetracycline (initiating) libraries from this dataset, using two replicates for each of the conditions (Supplementary Table S6). Model performance was evaluated with receiver operating characteristic (ROC) curves on the validation datasets for each replicate, the resulting AUC values of 0.9993 and 0.9994 in the tetracycline replicates were higher than those of chloramphenicol samples (0.9992 and 0.9983). The parameter importance in each of the models is shown in Supplementary Tables S7-10. The chloramphenicol models predicted a total of 3111 ORFs, including 57 elongations and 53 truncations (Supplementary Table S11). In the tetracycline dataset a total of 3711 ORFs were predicted, with 86 elongations and 79 truncations (Supplementary Table S12).

*E. coli* predictions were assessed against the ecogene curated set of 923 experimentally verified protein starts^33^. Genes within this dataset were excluded from the sets that were used to train the random forest models, in order to provide a means of assessing the accuracy of the ORF predictions. Five of the verified protein starts correspond to pseudogenes without annotated CDSs, of the remaining 917 verified protein starts, 821 (89.53%) matched ORFs in the tetracycline predictions, with 24 (2.62%) predicted ORFs in disagreement with the curated set (11 elongations, 13 truncations). In the chloramphenicol predictions 760 (82.88%) were found to match ecogene start sites, and 27 (2.94%) were found to be inconsistent (13 elongations, 14 truncations) with the verified protein starts (assuming genes do not have multiples TIS) (Supplementary Tables S11-12). Based on the experimentally verified starts the tetracycline-based classifier resulted in higher accuracy (sensitivity 0.9194, specificity 0.9996, positive predictive value 0.9716) than the chloramphenicol-based classifier (sensitivity 0.8539, specificity 0.9996, positive predictive value 0.9657). Surprisingly, the difference was not major arguing that using initiating ribosomes is not a requirement to obtain a good annotation of initiation sites.

## DISCUSSION

Our model shows that the distribution of ribo-seq footprint lengths can be used in conjunction with sequence features to accurately determine the translation initiation landscape of prokaryotes. These patterns are typically disrupted in standard ribo-seq analysis when reads of different fragment lengths are adjusted and merged to determine the specific codon under translation. The model is applicable across multiple organisms and experimental conditions and can be augmented with data from initiating ribosomes. It exhibits high accuracy as assessed by cross-validation, N-terminal proteomics and independent sequence-based metrics such as potential to form RNA structures. Interestingly, the predicted TISs exhibit known features of translation initiation which the previous annotations do not. In *S.* Typhimurium, our model provides evidence for 61 novel translated ORFs and the re-annotation of 419 genes. In particular, the current annotation includes 19 genes that lacked initiation codons, of which we were able to re-annotate 15 (Supplementary Table S4).

As expected, models based on initiating reads perform better than models based on elongating ribo-seq reads, suggesting that an optimal strategy for TIS identification would favour the use of the more focused, initiating ribo-seq profiles. However the degree of improvement between the models was relatively small, confirming the suitability of both elongating and initiating ribo-seq libraries for the purposes of TIS and ORF detection.

While the mechanistic or experimental origin of the patterns that our model captures remain unexplored, it is interesting to note the importance the models place (Supplementary Tables S1-2, S9-10) on the shorter range of fragments of 23-25 nt (*S*. Typhimurium) or 21-26 nt (*E. coli* tetracycline) in length. These shorter reads are consistent with recent reports of ribosomal subunits in a variety of distinct configurations, observed from translation complex profiling in the eukaryote *Saccharomyces cerevisiae*^34^. Whether similar patterns of read length distributions can be observed in eukaryotic ribo-seq datasets remains to be determined, although the method that we describe in this article is, regardless, fully extendable to eukaryotic datasets.

In conclusion, this study demonstrates the utility of ribo-seq fragment length patterns for TIS identification across multiple experimental conditions. These models provide a significant step forward in experimental TIS discovery, facilitating the move towards complete ORF annotation in both presumably well-annotated model organisms, as well as the ever growing list of newly sequenced genomes.

## ONLINE METHODS

### Preparation of ribo-seq libraries

Overnight stationary cultures of wild type *S.* Typhimurium (*Salmonella enterica* serovar Typhimurium - strain SL1344) grown in LB media at 37 °C with agitation (200 rpm) were diluted at 1:200 in LB and grown until they reached and OD600 of 0.5 (i.e., logarithmic (Log) phase grown cells). Bacterial cells were pre-treated for 5 min with chloramphenicol (Sigma Aldrich) at a final concentration of 100 μg/ml before collection by centrifugation (6000 × g, 5 min) at 4 °C. Collected cells were flash frozen in liquid nitrogen. The frozen pellet of a 50 ml culture was re-suspended and thawed in 1 ml ice-cold lysis buffer for polysome isolation (10 mM MgCl_2_, 100 mM NH_4_Cl, 20 mM Tris.HCl pH 8.0, 20 U/ml of RNase-free DNase I (NEB 2 U/μl), 1mM chloramphenicol (or 300 μg/ml), 20 μl/ml lysozyme (50mg/ml in water) and 100 u/ml SUPERase. In^™^ RNase Inhibitor (Thermo Fisher Scientific, Bremen, Germany)), vortexed and left on ice for 2 min with periodical agitation. Subsequently, the samples were subjected to mechanical disruption by two repetitive cycles of freeze-thawing in liquid nitrogen, added 5 mM CaCl_2_, 30μl 10% DOC and 1 × complete and EDTA-free protease inhibitor cocktail (Roche, Basel, Switzerland) and left on ice for 5 min. Lysates were clarified by centrifugation at 16,000 × g for 10 min at 4 °C.

For the monosome sample, the supernatant was subjected to MNase (Roche diagnostics Belgium) digestion using 600 U MNase (about ~1000 U per mg of protein). Digestion of polysomes proceeded for 1 h at 25 °C with gentle agitation at 400 rpm and the reaction was stopped by the addition of 10 mM EGTA. Next, monosomes were recovered by ultracentrifugation over a 1 M sucrose cushion in polysome isolation buffer without RNase-free DNase I and lysozyme, and with 2 mM DTT added using a TLA-120.2 rotor for 4 hr at 75,000 rpm and 4 °C.

For the selective purification of monosomes from polysomes (polysome sample), the supernatant was resolved on 10-55% (w/v) sucrose gradients by centrifugation using an SW41 rotor at 35,000 rpm for 2.5 hr at 4 °C. The sedimentation profiles were recorded at 260 nm and the gradient fractionated using a BioComp Gradient Master (BioComp) according to the manufacturer’s instructions. Polysome-enriched fractions were pooled and subjected to MNase digestion and monosome recovery as described above.

Ribosome-protected mRNA footprints with sizes ranging from 26-34 nt were selected and processed as described previously^14^ with some minor adjustments as previously described^35^. The resulting ribo-seq cDNA libraries of the monosome and polysome sample were duplexed and sequenced on a NextSeq 500 instrument (Illumina) to yield 75 bp single-end reads.

### Ribo-seq data processing

Ribo-seq data were preprocessed with cutadapt^36^ to remove sequencing adaptors, discarding reads less that 20 nt in length after trimming. Trimmed reads were initially aligned to the SILVA RNA database version 119^37^, the remaining reads were then mapped to either Salmonella enterica serovar Typhimurium - strain SL1344 (Assembly: GCA_000210855.2) or Escherichia coli str. K-12 substr. MG1655 (Assembly: GCA_000005845.2). Alignments were performed with bowtie2^38^. Reads were brought to codon resolution by adjusting the 5’ position of each read by a fixed distance offset, specific to each fragment length, based on visual identification of periodicity meta plots of the read counts per fragment length (Supplementary Figs. S5-6). In the *S.* Typhimurium dataset the following fragments lengths were selected and adjusted by the values in brackets, in the monosome sample 29 (13 nt), 30 (14 nt), 31 (15 nt), 32 (16 nt), 33 (17 nt), and in the polysome sample 29 (13 nt), 30 (14 nt), 31 (15 nt), 32 (16 nt), 33 (17 nt) and 34 (18 nt). Selected reads of the indicated lengths account for 39.98 and 48.69 % of total reads for the monosome and polysome samples, respectively.

Recent publications reporting prokaryotic ribo-seq^26,28,39,40^ suggest that read fragments from libraries digested with micrococcal nuclease align more precisely to their 3’ rather than 5’ ends. Consistent with this, we observe a modest increase in the periodicity of meta profiles of the *S.* Typhimurium ribo-seq libraries when reads are brought to codon resolution from the 3’ end (Supplementary Fig. S1), however this does not hold true for the *E. coli* datasets, where the use of 3’ poly adenosine adaptors, results in a loss of resolution at the 3’ end after read trimming (Supplementary Fig. S1), making the use of 5’ ends preferable.

Regardless, the protected read fragment patterns that we use in the input feature vectors for the classifier takes both length and position into consideration. Consequently the classifier is unaffected by this choice. However, to maintain consistency throughout this study read counting for model predictors was performed from the 5’ end for all libraries.

### Read distributions and heat maps

Ribo-seq read distributions were summarised over all annotated start codons in the *S.* Typhimurium and *E. coli* annotations respectively. 5’ read counts were taken from regions 30nt upstream to 60nt downstream of the start codon, 3’ read counts were taken from the first nucleotide of the start codon up to 90 nucleotides downstream. All reads with a MAPQ greater than 10, from the upper 90% of genes by total CDS expression were included. Total counts were scaled to a sum of one per individual region, in order to not disproportionately favour profiles from highly expressed genes. Meta plots were then produced to show the proportion of read counts over the window across all genes. 3’ and 5’ heatmaps were generated from the scaled regions, showing the number of standard deviations from the row (fragment length) mean.

### Model implementation

For each candidate TIS a feature vector was defined as each nucleotide in a -20 to +10nt window around the position, the ribo-seq 5’ FPKM (fragments per kilobase per million mapped reads) between the current position and the next in-frame downstream stop codon, the count of potential in-frame start sites (codons within one edit distance of ATG) from the nearest in-frame upstream stop codon to the current position, the proportion of 5’ ribo-seq reads upstream in a 20 nt window, the proportion of 5’ ribo-seq reads downstream in a 20 nt window, the ratio of 5’ ribo-seq reads up and downstream and the proportion 5’ ribo-seq counts per fragment length for a fixed range of positions in relation to current site (selected from visual inspection of 5’ fragment length heatmaps (Supplementary Fig. S4). In the *S.* Typhimurium samples fragment lengths 20-35 nt in positions -20 to -11 and 0 nt, were used. In the *E. coli* datasets for the tetracycline samples fragment lengths 20-35 nt at positions -25 to -16 nt, were selected and for chloramphenicol lengths 30 to 50 nt, at positions -25 to -16 and -1 to +1nt were used.

Stop-to-stop windows were defined for each annotated gene as all in-frame positions between the nearest in-frame upstream stop codon and the stop codon of the gene (with a maximum length cut-off 999 nt upstream).

The H2o random forest implementation^41^ was used and the models were trained with positive examples of randomly selected annotated start codons from the upper 50% of genes ranked by ribo-seq expression over the gene CDS. We additionally required that the positive examples were not among the genes supported by N-terminal peptides in the *S.* Typhimurium samples or included in the ecogene dataset for the *E. coli* samples, since these were retained for model accuracy assessment. Negative examples were randomly selected from in-frame codons in the stop-to-stop windows both upstream and downstream of the annotated TIS. The *S.* Typhimurium models were trained on 1500 positive and 6000 negative positions, with an independent validation set of 200 positive and 800 negative positions, the validation set was used for parameter training (number of trees monosome: 600, polysome: 600). The *E. coli* models were trained on 1100 positive and 4400 negative positions, with an independent validation set of 200 positive and 800 negative positions for parameter tuning (number of trees: CM1: 950, CM2: 950, TET2: 650 and TET3: 700). Predictions were then run against all cognate and near cognate (defined as one edit distance from ATG) in-frame positions, in the stop-to-stop regions. Novel predictions were performed against all cognate and near cognate codons in stop-to-stop regions around ORFs of at least 300nt in length, with a ribo-seq read coverage of 0.75 or more (ORF coverage was defined as the proportion of nucleotides in each predicted ORF that at least one ribo-seq read mapped to), that did not overlap with annotated exons. ORFs were delineated by extending each candidate TIS to the closest in-frame stop codon. For a given stop-to-stop region the model selected the TIS with the highest positive predicted score per sample. Predictions from the replicates for each of the datasets were then compared, discarding predictions that were unique to only one replicate.

### N-terminal proteomics

Overnight stationary cultures of wild type *S.* Typhimurium (*Salmonella enterica* serovar Typhimurium - strain SL1344) grown in LB media at 37 °C with agitation (200 rpm) were diluted at 1:200 in LB and grown until they reached and OD600 of 0.5 (i.e., logarithmic (Log) phase grown cells). Bacterial cells were collected by centrifugation (6000 × g, 5 min) at 4 °C, flash frozen in liquid nitrogen and cryogenically pulverized using a liquid nitrogen cooled pestle and mortar. The frozen pellet of a 50 ml culture was re-suspended and thawed in 1 ml ice-cold lysis buffer (50 mm NH_4_HCO_3_ (pH 7.9) supplemented with a complete protease inhibitor cocktail tablet (Roche Diagnostics GmbH, Mannheim, Germany) and subjected to mechanical disruption by two repetitive freeze-thaw and sonication cycles (i.e. 2 minutes of sonication on ice for 20-s bursts at output level 4 with a 40% duty cycle (Branson Sonifier 250; Ultrasonic Convertor)). The lysate was cleared by centrifugation for 15 min at 16,000 × *g* and the protein concentration measured using the protein assay kit (Bio-Rad) according to the manufacturer’s instructions. The lysate was added Gu.HCl (4M f.c.) and subjected to N-terminal COFRADIC analysis as described previously^42^. Free amines were blocked at the protein level making use of an N-hydroxysuccinimide ester of (stable isotopic encoded) acetate (i.e. NHS esters of ^13^C_2_D_3_ acetate), which allows distinguishing in vivo and in vitro blocked N-terminal peptides^43^. The modified protein sample was digested overnight with sequencing-grade modified trypsin (1/100 (w/w trypsin /substrate)) at 37 °C and subsequent steps of the N-terminal COFRADIC procedure were performed as previously described^42^.

### LC-MS/MS analysis

LC-MS/MS analysis was performed using an Ultimate 3000 RSLC nano HPLC (Dionex, Amsterdam, the Netherlands) in-line connected to an LTQ Orbitrap Velos mass spectrometer (Thermo Fisher Scientific, Bremen, Germany). The sample mixture was loaded on a trapping column (made in-house, 100 μm I.D. × 20 mm, 5 μm beads C18 Reprosil-HD, Dr. Maisch). After back flushing from the trapping column, the sample was loaded on a reverse-phase column (made in-house, 75 m I.D. × 150 mm, 5 μm beads C18 Reprosil-HD, Dr. Maisch). Peptides were loaded in solvent A’ (0.1% trifluoroacetic acid, 2% acetonitrile (ACN)) and separated with a linear gradient from 2% solvent A’’ (0.1% formic acid) to 50% solvent B’ (0.1% formic acid and 80% ACN) at a flow rate of 300 nl/min followed by a wash reaching 100% solvent B’. The mass spectrometer was operated in data-dependent mode, automatically switching between MS and MS/MS acquisition for the ten most abundant peaks in a given MS spectrum. Full scan MS spectra were acquired in the Orbitrap at a target value of 1E6 with a resolution of 60,000. The 10 most intense ions were then isolated for fragmentation in the linear ion trap, with a dynamic exclusion of 20 s. Peptides were fragmented after filling the ion trap at a target value of 1E4 ion counts.

Mascot Generic Files were created from the MS/MS data in each LC run using the Mascot Distiller software (version 2.5.1.0, Matrix Science, www.matrixscience.com/Distiller.html). To generate these MS/MS peak lists, grouping of spectra was allowed with a maximum intermediate retention time of 30 s and a maximum intermediate scan count of 5. Grouping was done with a 0.005 Da precursor tolerance. A peak list was only generated when the MS/MS spectrum contained more than 10 peaks. There was no de-isotoping and the relative signal-to-noise limit was set at 2.

The generated MS/MS peak lists were searched with Mascot using the Mascot Daemon interface (version 2.5.1, Matrix Science). Searches were performed using a 6-FT database of the *S.* Typhimurium genome combined with the Ensembl protein sequence database (assembly AMS21085v2 version 86.1), which totalled 139,408 entries after removal of redundant sequences. The 6-FT database was generated by traversing the entire genome across the six reading frames and searching for all NTG (N=A,T,C,G) start codons and extending each to the nearest in frame stop codon (TAA,TGA,TAG), discarding ORFs less than 21nt in length. The Mascot search parameters were set as follows: Heavy acetylation at lysine side-chains (Acetyl:2H(3)C13(2) (K)), carbamidomethylation of cysteine and methionine oxidation to methionine-sulfoxide were set as fixed modifications. Variable modifications were formylation, acetylation and heavy acetylation of N-termini (Acetyl:2H(3)C13(2) (N-term)) and pyroglutamate formation of N-terminal glutamine (both at peptide level). Endoproteinase semi-Arg-C/P (semi Arg-C specificity with Arg-Pro cleavage allowed) was set as enzyme allowing for no missed cleavages. Mass tolerance was set to 10 ppm on the precursor ion and to 0.5 Da on fragment ions. Peptide charge was set to 1 +, 2+, 3+ and instrument setting was put to ESI-TRAP. Only peptides that were ranked one, have a minimum amino acid length of seven, scored above the threshold score, set at 95% confidence, and belonged to the category of *in vivo* or *in vitro* blocked N-terminal peptides compliant with the rules of initiator methionine (iMet) processing^44^ were withheld. More specifically, iMet processing was considered in the case of iMet-starting N-termini followed by any of the following amino acids; Ala, Cys, Gly, Pro, Ser, Thr, Met or Val and only if the iMet was encoded by ATG or any of the following near-cognate start codons; GTG and TTG (Supplementary Table S13). In contrast to eukaryotic nascent protein N-termini, the typical lack of significant steady-state levels of N-terminal protein modification (e.g. Nt-acetylation or Nt-formylation), warrant caution to unequivocally assign bacterial protein N-termini as proxies of translation initiation. As such, a high confidence subset of Nt-formylated Met-starting N-termini, was selected (Supplementary Table S14).

### Assessing model accuracy

GC content was calculated at the third nucleotide positions for all annotated and predicted ORFs and mean GC values were summarised for each subgroup of predicted ORFs (matching annotations, truncations and elongations) in 9 nt sliding windows, over regions 57 nt upstream and 57 nt downstream of the annotated or predicted start sites.

-20 to + 20 nt nucleotide sequences were extracted around the predicted and annotated TIS in the *S.* Typhimurium and *E. coli* genomes. Sequence logos were generated for each subgroup of matching annotations, truncations, elongations and novel genes, using the weblogo tool^45^.

The minimum free energy of RNA secondary structure around predicted and annotated ORFs was estimated with RNAfold version 2.1.9 from the ViennaRNA package^46^. Mean free energy values were summarised for each ORF class in 39 nt sliding window across regions 57 nt up and downstream of the start codon.

Read distributions were created for each subgroup of predicted ORFs (matching annotations, truncations, elongations, and novel genes) and their corresponding annotated TIS. Distributions of ribo-seq reads adjusted to codon level resolution, were summarised in regions 30 nt upstream and downstream of the first nucleotide of the initiation codon, total counts of each individual region were scaled to a sum of one, in order to normalise profiles for differences in gene expression levels. Meta plots were then produced to show the proportion of reads over the window position from all predicted subgroups and their corresponding annotated start codons.

Sensitivity, specificity and positive predictive values were calculated from all genes that were supported by either high confidence n-terminal peptides (*S*. Typhimurium) or experimentally verified protein starts (*E. coli).* Supported predicted ORFs were considered true positives, predicted ORFs that disagreed with supported positions were classified as false positives. False negatives were assigned from supported genes where no ORF was predicted. All in-frame cognate and near cognate start codons (one edit distance from ATG), in CDS regions of supported genes that were neither predicted nor supported were considered true negatives.

Amino acid sequences of novel ORF were compared to known proteins in the nonredundant protein database (Update date:2016/12/15) and protein domains (cdd.v.3.15) using BLASTP^47^ (version 2.5.1+). Hits with the greatest coverage of query sequence and lowest e-value were selected. Hits were considered highly similar if they shared >95% identify to a protein sequence, over 100% of the novel ORF sequence

## DATA AVAILABILITY

The previously published *E. coli* ribo-seq dataset was downloaded from the NCBI SRA (BioProject ID:PRJDB2960). *S.* Typhimurium ribo-seq sequencing data has been deposited in NCBI’s Gene Expression Omnibus^48^ and is accessible through GEO Series accession number GSE91066. *S.* Typhimurium mass spectrometry proteomics data have been deposited to the ProteomeXchange Consortium via the PRIDE^49^ partner repository with the dataset identifier PXD005579 and 10.6019/PXD005579.

## ACKNOWLEDGMENTS

We thank Gunnar Schulze and Kornel Labun for valuable discussions. Prof. Kris Gevaert for financial support of this research (Research Foundation - Flanders (FWO-Vlaanderen), project number G.0440.10). P.V.D. acknowledges support from the Research Foundation - Flanders (FWO-Vlaanderen), project number G.0269.13N. A.G and E.V. acknowledge support from the Bergen Research Foundation.

## AUTHOR CONTRIBUTIONS

A.G., E.V. and P.V.D. conceived the study and wrote the manuscript; A.G. performed the computational analysis; P.V.D performed the proteomics experiment. E.N. and P.V.D. performed proteomics analysis; P.V.D. and V.J. prepared the ribo-seq libraries.

